# G-quadruplex RNA motifs influence gene expression in the malaria parasite *Plasmodium falciparum*

**DOI:** 10.1101/2021.03.08.434398

**Authors:** Franck Dumetz, Eugene Yui-Ching Chow, Lynne M. Harris, Mubarak I. Umar, Anders Jensen, Betty Chung, Ting Fung Chan, Catherine J. Merrick, Chun Kit Kwok

## Abstract

G-quadruplexes are non-helical secondary structures that can fold *in vivo* in both DNA and RNA. In human cells, they can influence replication, transcription and telomere maintenance in DNA, or translation, transcript processing and stability of RNA. We have previously showed that G-quadruplexes are detectable in the DNA of the malaria parasite *Plasmodium falciparum*, despite a very highly A/T-biased genome with unusually few guanine-rich sequences. Here, we show that RNA G-quadruplexes can also form in *P. falciparum* RNA, using rG4-seq for transcriptome-wide structure-specific RNA probing. Many of the motifs, detected here via the rG4seeker pipeline, have non-canonical forms and would not be predicted by standard *in silico* algorithms. However, *in vitro* biophysical assays verified the formation of non-canonical motifs. The G-quadruplexes in the *P. falciparum* transcriptome are frequently clustered in certain genes and associated with regions encoding low-complexity peptide repeats. They are overrepresented in particular classes of genes, notably those that encode *Pf*EMP1 virulence factors, stress response genes and DNA binding proteins. *In vitro* translation experiments and *in vivo* measures of translation efficiency showed that G-quadruplexes can influence the translation of *P. falciparum* mRNAs. Thus, the G-quadruplex is a novel player in post-transcriptional regulation of gene expression in this major human pathogen.

## INTRODUCTION

Protozoan *Plasmodium* parasites are the causative agents of human malaria, a disease responsible for widespread morbidity and almost half a million deaths each year ^1^. Most of the deaths are caused by the species *P. falciparum*, which has an unusual genome with an extreme A/T-bias of ~81% A/T ^2^. Not all *Plasmodium* species share this feature: the five other *Plasmodium* species that infect humans have genomes with A/T contents of ~60%, 61%, 71% and 76% for *P. vivax, P. knowlesi, P. ovale curtisi/wallikeri* and *P. malariae* respectively ^3^. In *P. falciparum*, however, the particularly A/T-biased genome results in an extreme paucity of guanine-rich sequences, and hence of putative G-quadruplex forming sequences (PQSs) ^4,5^.

The G-quadruplex is an important non-double-helical structure that can form in DNA and also RNA ^6^. It is classically formed from four tracts of at least three guanines found in close proximity on the same strand: these are arranged as three or more quartets stacked on top of one another ^7^. The requisite guanine density is a rare feature in A/T-biased genomes, but the existence of G-quadruplexes has nevertheless been predicted in the *P. falciparum* genome by searching for the consensus sequence (G_3_ N_x_)_4_ ^4,5,8^. Only ~100 PQSs are found outside the inherently-G-rich telomeres: this is approximately one per 300kb of the non-telomeric genome, compared to an average of one per kb in the human genome ^9^. More modern algorithms such as *G4Hunter* ^10^, which can identify PQSs in ‘non-canonical’ forms beyond the classical (G_3_ N_x_)_4_, can increase the PQS count in the *P. falciparum* genome, but only modestly ^5,11^.

Despite their scarcity, some G-quadruplexes have been shown to fold in *P. falciparum* DNA both *in vivo* ^12^ and *in vitro* ^5,8,11–13^. By contrast, RNA G-quadruplexes (rG4s) have to date remained unexplored in *P. falciparum*, or in any other organism with a comparable genome bias. Nevertheless, they could have important roles in parasite biology. PQSs in the *P. falciparum* genome are primarily found around the major virulence gene family *var* – either within the coding sequence or upstream of *var* genes ^4,5,8^. Mitotic recombination amongst these virulence genes is strongly spatially associated with PQSs ^4^, and the loss of RecQ helicases, which normally act to unwind G-quadruplexes, can increase the rate of *var* gene recombination ^14^. Therefore, one unique role for G-quadruplexes at the DNA level in *P. falciparum* is probably to promote *var* gene diversification. If these G-quadruplexes also occurred at the RNA level in *var* gene transcripts, they could modulate expression at the translational level. Similarly, translation could be affected in non-*var* genes that harbour rG4s. It has been suggested that post-transcriptional regulation might be particularly important in *P. falciparum* because this organism has unusually low numbers of sequence-specific transcription factors ^15,16^ and a ‘hardwired’ transcriptional cascade across its cell cycle ^17^.

In several model organisms, the rG4 content of the transcriptome has recently been investigated by ‘rG4-seq’ ^18–20^, a technique in which the transcriptome is extracted from a cell and reverse-transcribed in two parallel conditions, one allowing rG4s to fold and the other preventing them from folding. Folded rG4s tend to stall the reverse transcriptase, leading to rG4-specific truncations that can be detected by comparing the two resultant cDNA libraries. This method has detected many rG4s in the transcriptomes of human, plant and bacterial cells ^18–20^, some of which have functional consequences in gene translation ^19–22^. Here we performed rG4-seq on the highly A/U-biased transcriptome of *P. falciparum* and investigated the biological roles of rG4s in this parasite.

## RESULTS

### G-quadruplexes can be detected in *P. falciparum* RNA

The most direct route to demonstrate the presence of G-quadruplexes in cells is arguably to visualise them with microscopy. Cell-permeable G-quadruplex-specific fluorescent dyes have therefore been developed, some of which are specific for RNA rather than DNA ^23^. Structure-specific antibodies also exist ^6^ but do not distinguish RNA and DNA, although some antibodies can detect both structures ^24^.

We attempted to visualise rG4s specifically in cultured *P. falciparum* parasites using the rG4 dye QUMA-1 ^23^ (Supplementary Figure 1). A fluorescent signal was seen throughout the parasites and this appeared to be somewhat RNase-sensitive, but much of the signal was apparently nuclear and DNase-sensitive, suggesting that the QUMA-1 dye is not as RNA-specific in *P. falciparum* cells as it is in human cells DNA G4s are likely to dominate in parasite nuclei, particularly in telomeres, while rG4s, if present, are likely to be less abundant and more dispersed throughout the cytoplasm, so they may be difficult to detect via microscopy.

We proceeded to use rG4-seq ^18^ to detect rG4s and to obtain a comprehensive catalogue of these motifs in the *P. falciparum* transcriptome. We conducted rG4-seq using RNA extracts obtained from duplicate cultures of mixed-stage asexual parasites (3D7 strain), enriched for polyadenylated RNA. Under an *in vitro* rG4 stabilizing condition (buffer containing 100 mM K^+^), approximately 300 reverse transcriptase stalling (RTS) events associated with PQSs were detected by rG4-seq (Figure 1A). The number of detections further increased to more than 2000 when the rG4 stabilizing ligand pyridostatin (PDS) was included in the buffer (Figure 1A). Across all four rG4-seq experiments (2 replicates, 2 conditions), >95% of the RTS events captured were associated with various categories of PQS, which confirmed that these events were highly specific to genuine rG4s and that the false discovery rate was low (Figure 1A). By comparing the detection results between biological replicates, we found that >70% of the RTS events were reproduced, coherent with the outcomes from an earlier rG4-seq experiment in human cells ^18^ (Figure 1B, C). Reproducibility was best amongst what might be considered the highest-confidence PQSs: both canonical and non-canonical rG4 motifs (the non-canonical category including bulged and 2-quartet motifs) (Figure 1B). Reproducibility was lower amongst the set of guanine-rich sequences that may have G4-forming potential, but with structural imperfections (Figure 1C). Overall, the experiments confirmed that rG4s can fold in *P. falciparum* mRNAs, with a unified set of 2,569 rG4s detected (Supplementary Datasheet 1).

**Figure 1:**
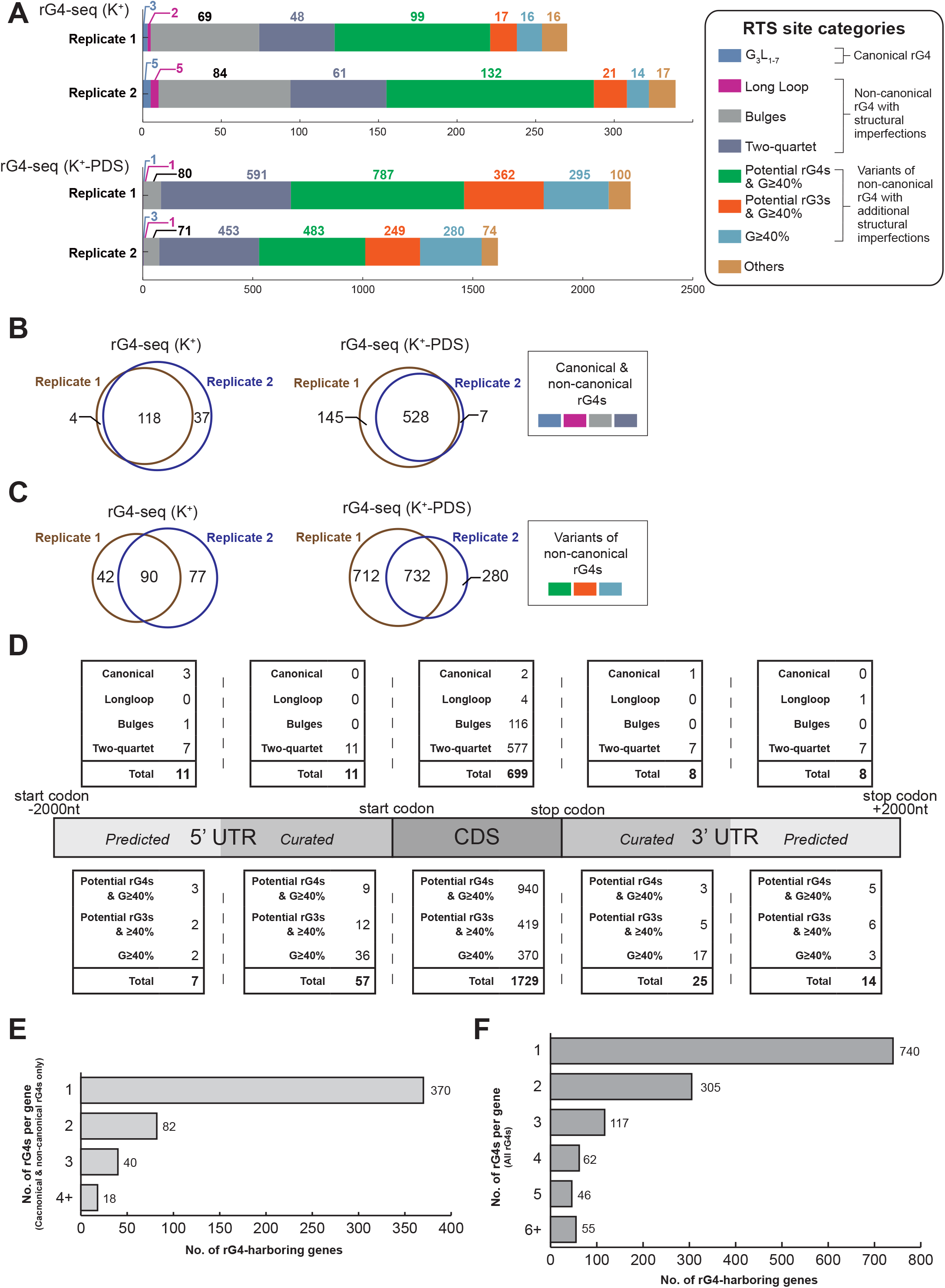
The landscape of rG4s in the *P. falciparum* transcriptome. (A) Distribution and reproducibility of RTS sites detected in rG4-seq experiments. RTS sites were assigned to rG4 structural classes according to their adjacent nucleotide sequences. RTS sites classified as “Others” were considered false positive detections as their adjacent nucleotide sequences do not satisfy the minimum requirements for forming quadruplex or triplex structures. (B,C) Reproducibility of RTS sites detected by rG4-seq experiments for the rG4s in the four well-established structural classes (B) and in other classes (C). K^+^ refers to the rG4-seq experiments conducted in potassium ions alone; K^+^-PDS refers to RNA folding in potassium plus pyridostatin. (D) Distribution of detected rG4s of seven different classes within the coding regions and untranslated regions of *P. falciparum* protein coding genes. (E,F) Summary of the number of rG4s detected per *P. falciparum* gene, considering only canonical and non-canonical rG4 motifs (E) or all rG4 motifs (F).

### Detected RNA G-quadruplexes in *P. falciparum* are mostly non-canonical

The detected rG4s in the *P. falciparum* transcriptome were mostly located in the CDS regions of protein coding genes, rather than in the UTRs (Figure 1D). Moreover, based on their nucleotide sequences, most of the rG4s were classified as non-canonical motifs (e.g. bulged or two-quartet) (Figure 1A, D), similar to the findings of rG4-seq in human cells ^18^. These imperfections do not necessarily compromise rG4 folding but they have been suggested to have a negative impact on the thermostability of the resultant rG4 structures ^18,25^. The *P. falciparum* transcriptome also harboured many guanine-rich potential rG4s and RNA G-triplex (rG3) motifs that could potentially fold into quadruplex/triplex structures despite their nucleotide sequences not matching the four better-established structural motif definitions ^26^.

The rG4-seq results appeared to contrast with previous *in silico* analyses of the *P. falciparum* genome for PQSs that might form in DNA ^4,8^: those analyses examined only PQSs of the form (G_3_N_x_)_4_ and ~100 were found, most notably among the *var* virulence genes that encode *P. falciparum* Erythrocyte Membrane Protein 1 (*Pf*EMP1). The rG4-seq experiment revealed very few rG4s of canonical or long-loop motif types – either in *var* genes or elsewhere in the transcriptome (Figure 1A, D). Nevertheless, rG4s appeared overall to be quite prevalent among *P. falciparum* genes: out of 5,762 annotated genes, nearly 10% of them (510 genes) contained at least one rG4 from the four better-established structural motifs (Figure 1E) and this rose to 22% (1,325 genes) if the potential rG4/rG3 structures and G≥40% motifs were also considered (Figure 1F). It was also not uncommon to find genes harbouring two or more rG4s simultaneously (Figure 1E, F). Given the low overall G/C content of the *P. falciparum* transcriptome, the prevalence of rG4s in these genes is striking.

### The *P. falciparum* transcriptome does contain some canonical rG4s

To further explore any perceived discrepancies between rG4-seq results and prior *in silico* analyses of the *P. falciparum* genome ^4,8^, we first updated the *in silico* PQS prediction on the *P. falciparum* transcriptome for canonical or long-loop structural motifs fitting the (G_3_N_x_)_4_ criteria (Supplementary Datasheet 2). The predicted PQSs were then further curated for any overlap with DNA repetitive elements using RepeatMasker software and/or manual curation (Supplementary Datasheet 2). Among the predicted 355 PQSs across 41 genes, it was striking that the vast majority of them overlapped with telomeric GGGTT(C/T)A repeats, AAAGGGG repeats, or regions of exact homology within genes, and that none of these was detected by rG4-seq (Supplementary Table 1, Figure 2A). In contrast, among the 28 predicted PQSs that did not overlap with repetitive elements, 12 of them were detected by rG4-seq (Supplementary Table 1). This suggested that the technique can only detect predicted PQSs in the *P. falciparum* transcriptome if they do not overlap with repetitive elements – probably due to technical limitations in unambiguously assigning short-read Illumina sequencing data to repetitive sequences. Although our results cannot offer reliable information on PQSs associated with repetitive sequences, we can still confirm that: a) outside of repetitive regions, canonical and long-loop rG4s are scarce in the *P. falciparum* transcriptome, and b) virulence genes including *var* and *rifin* (as well as a small number of other genes), do harbour a few non-repeat-associated canonical/long-loop rG4s in their CDS and/or UTRs.

**Figure 2:**
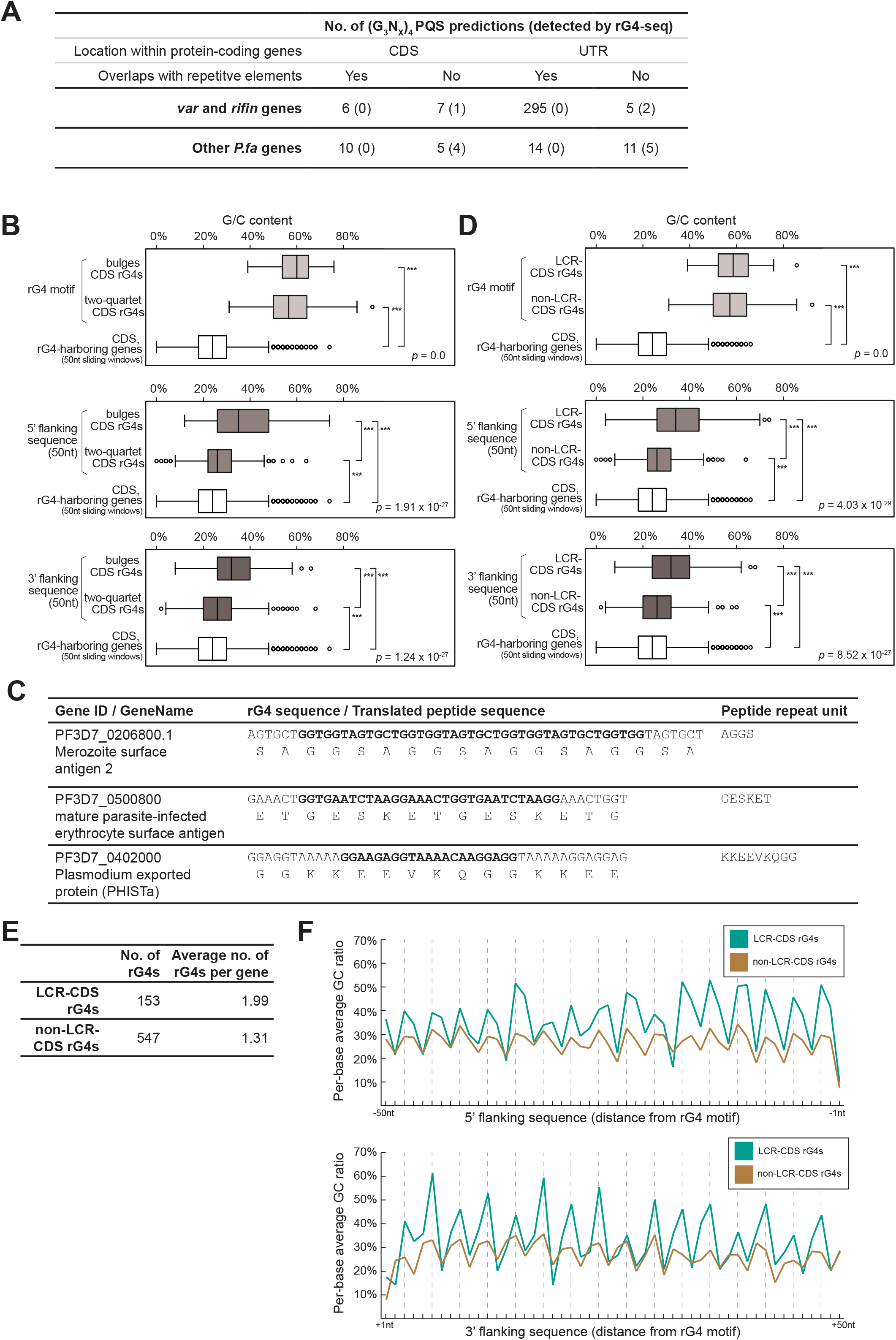
Sequence features of *P. falciparum* RNA G-quadruplexes. (A) Summary of (G_3_N_x_)_4_ PQS prediction outcomes in protein-coding genes and their status of detection in rG4-seq. PQSs matching the canonical or long-loop motif definitions were considered as (G_3_N_x_)_4_ PQSs. Their location and overlapping status with repetitive elements are highlighted. (B) Comparative analysis of G/C content in CDS rG4s categorised as either bulged or two-quartet structural motifs. The G/C content of the rG4 motif, the rG4 flanking regions and the whole rG4-harbouring gene are plotted. P-values shown in each subplot indicates the outcome of a Kruskal– Wallis one-way ANOVA test comparing the 3 samples. *** indicates pairwise statistical significance in the difference between G/C contents (*p* <0.005, Wilcoxon rank-sum test); ns indicates not significant. (C) Examples of CDS rG4s that overlap with gene sequences encoding low-complexity peptide regions. (D) Comparative analysis of G/C content in CDS rG4s that are adjacent/not-adjacent to low-complexity peptide regions (LCRs), regarding the G/C content of the rG4 motif, the rG4 flanking regions and the whole rG4-harbouring gene. P-values shown in each subplot indicates the outcome of a Kruskal–Wallis one-way ANOVA test comparing the 3 samples. *** indicates pairwise statistical significance in the difference between G/C contents (*p* <0.005, Wilcoxon rank-sum test); ns indicates not significant. (E) Summary of the numbers and the degree of clustering of CDS rG4s that are adjacent/not-adjacent to LCRs. (F) Comparison of per-base average G/C content in the flanking regions of LCR CDS rG4s and non-LCR CDS rG4s.

### Detected RNA G-quadruplexes are non-randomly distributed in the *P. falciparum* transcriptome

We next explored the distribution and sequence characteristics of non-canonical CDS rG4s in the bulged and two-quartet categories. These formed the bulk of the detected rG4s (and were not considered in previous analyses focussing on DNA PQSs because such motifs are not expected to be stable in DNA). We calculated the G/C content of each rG4 sequence and its 50nt 5’/3’ flanking sequences, then compared these with the G/C content of the CDS of the whole rG4-harbouring gene in 50nt sliding windows. As expected, the G/C content was highly elevated in the rG4 sequences themselves (Figure 2B). In addition, the flanking regions of bulged rG4 motifs had significantly higher G/C contents than the rest of the transcript, whereas this difference was much more subtle in the flanking regions of two-quartet rG4 motifs (Figure 2B).

Through manually examining the nucleotide sequences of bulged rG4 motifs and their translated peptide sequences, we found that some of these rG4s overlapped with low-complexity peptide regions (LCRs) with a compositional bias of amino acid residues and/or tandem repeats of short peptides (Figure 2C). This observation motivated us to search *de novo* for LCRs in the codons that encoded detected rG4s, and then to annotate all non-canonical CDS rG4s based on their association with LCRs. Strikingly, we found that rG4s located in LCRs possessed distinct features. They were associated with a higher G/C content in the flanking regions (Figure 2D) and with multiple occurrences of rG4s within a single gene (Figure 2E).

We also examined the per-base G/C content of 5’ and 3’ flanking regions and observed a three-base periodicity in the average G/C content, reflecting the property of protein-coding sequences (Figure 2F). Furthermore, this revealed that the G/C difference between flanking regions of LCR and non-LCR rG4s was driven by the strength of the G/C spikes occurring once every three bases (Figure 2F). This important observation suggests that common repetitive elements in the *P. falciparum* genome, such as the GGGTTTA telomeric repeat and other repeats, are unlikely to be a direct cause of LCR CDS rG4s, since their repeat units are usually 7/8nts long and not a multiple of 3.

### RNA G-quadruplexes tend to occur in genes with specific GO terms

To evaluate the potential functional significance of *P. falciparum* rG4s, we explored the associations between rG4-harbouring genes and *Plasmodium* biology via a gene ontology (GO) term enrichment analysis conducted on the rG4-harbouring genes. Some enriched GO terms appeared, but they were very broad and covered too many genes to define a clear biological process (Figure 3A). We therefore altered the analysis strategy to identify the most common GO terms supported by five or more rG4-harbouring genes (Figure 3B). This revealed at least three categories of interest: firstly, there were multiple GO terms related to pathogenesis, including antigenic variation and cell-cell adhesion – terms that primarily apply to variantly-expressed virulence genes such as *var* genes, confirming the earlier finding that *var* and *rifin* genes can encode rG4s. Secondly, there were multiple terms relating to stress-responsiveness, including proteotoxic stress and oxidative stress, both of which can be induced by human immune responses to malaria and by antimalarial drugs – indeed, the term ‘response to drug’ appeared as well. Thirdly, genes involved in transcription and regulation of transcription such as ApiAP2 transcription factors and chromatin modifiers were enriched: this is consonant with the rG4-seq results from human cells, in which many genes encoding DNA-binding and DNA-modifying proteins were detected ^18^.

**Figure 3:**
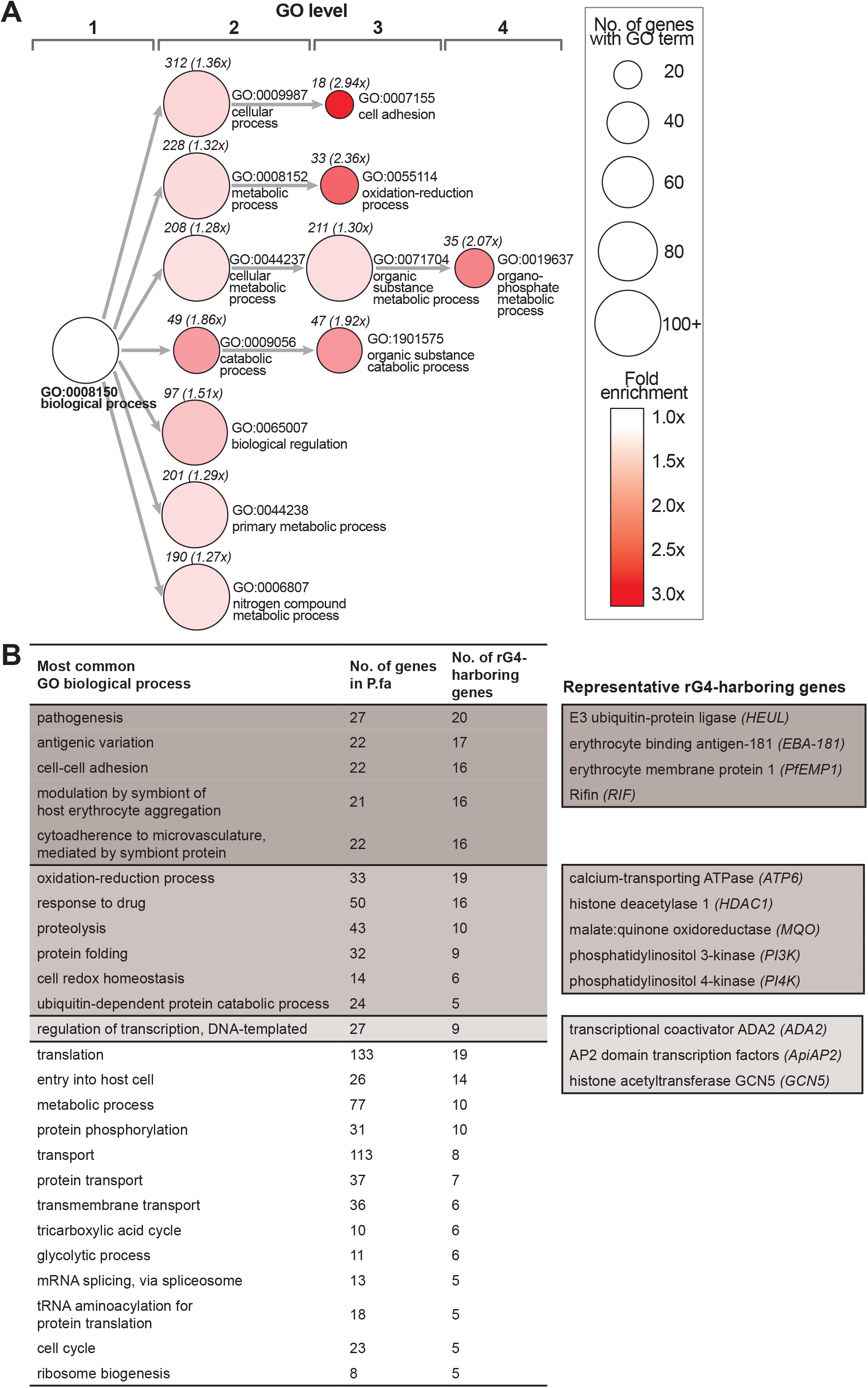
Functional associations of rG4-harbouring protein-coding genes. (A) Results of GO ‘biological process’ enrichment analysis on genes that harbour rG4s belonging to the 4 canonical/non-canonical rG4 structural motifs. Only enriched GO terms with a p value < 0.05 and FDR < 0.05 are shown. (B) Summary of the most common GO biological processes among genes that harbour rG4s belonging to the 4 canonical/non-canonical rG4 structural motifs. Three groups of biologically-related GO terms are highlighted, with examples of corresponding rG4-harbouring genes.

### An RNA G-quadruplex can affect *in vitro* translation of a *P. falciparum* gene

Having analysed the rG4-seq dataset *in silico*, we proceeded to investigate the biological effects of *Plasmodium* rG4s via *in vitro* and *in vivo* experiments. To assess the potential effect of rG4s on *P. falciparum* protein production we first conducted *in vitro* translation experiments. The gene PF3D7_0934400, encoding an ApiAP2 protein, was selected: it has a 2-quartet PQS on the coding strand which was found as an rG4 in our dataset. A second gene, PF3D7_1254400, was selected as a control: this encodes a rifin protein and has a PQS on the non-coding strand ^4^, so it was not found in our rG4 dataset. Both genes were cloned into an *in vitro* expression vector, pTnT-HiBiT, containing a T7 RNA polymerase promotor and a HiBiT bioluminescent tag. The original gene sequences were termed wild-type (WT) and site-directed mutagenesis was then used to derive ‘G4mut’ constructs wherein the PQS was disrupted (Figure 4A).

**Figure 4:**
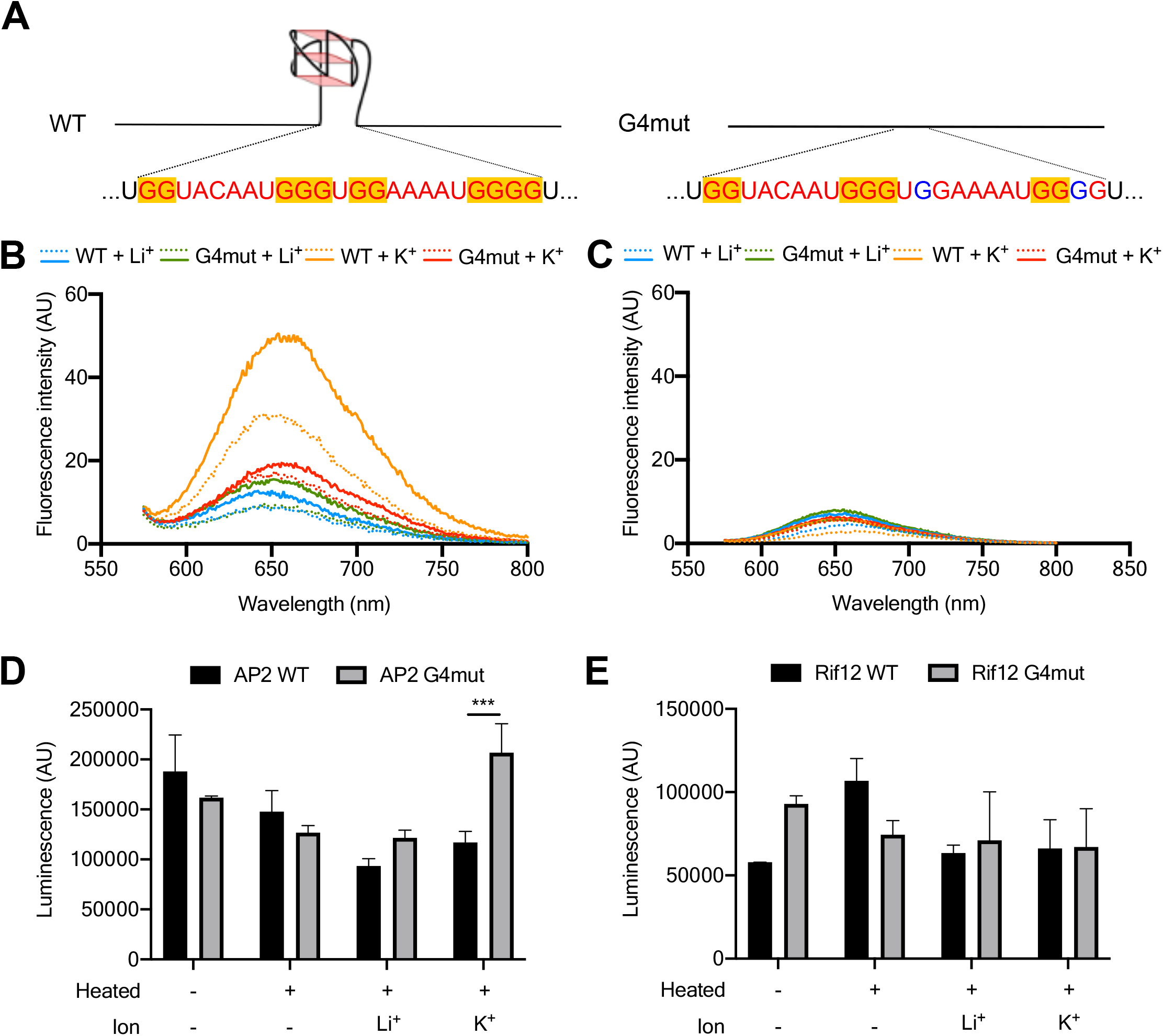
*In vitro* folding of rG4 motifs in reporter genes and their impact on *in vitro* translation. (A) Schematic showing the WT and G4mut forms of the *AP2* gene with its rG4-forming motif intact or mutated. (B,C) QUMA-1 fluorescence assay respectively on AP2 (B) and Rifin12 (C) rG4-only oligos (dashed line) and full-length transcripts (solid line) in the presence of LiCl or KCl. Two versions of each transcript were tested, the WT transcript and the rG4mut transcript where the sequence has been modified to prevent rG4 formation. (D,E) *In vitro* translation of *AP2* and *Rifin*12 using the nano-luciferase HiBiT moiety as the detection method. The previously folded transcripts were used on rabbit reticulocyte lysate to produce the protein.

The formation of the rG4 in the *ApiAP2* gene was confirmed using four different biophysical experiments (Supplementary Figure 2) conducted on WT and G4mut RNA oligos (Supplementary Table 2). We also assessed rG4 formation in full-length transcripts after *in vitro* translation – the first time, to our knowledge, that such an assessment has been made in full-length transcripts. For this, we used QUMA-1 as the rG4-specific binding agent ^23^. QUMA-1 fluorescence was much higher from the WT gene (both full-length and RNA oligo) than from the G4mut form (Figure 4B) and this required a G4-stabilising cation, i.e. K^+^ rather than Li^+^. The same experiment performed on the *rifin* gene (both full-length transcript and RNA oligos) gave no enhanced QUMA-1 fluorescence (Figure 4C), consistent with the absence of rG4s. Collectively, the results indicate the formation of an rG4 in the *ApiAP2* WT transcript and not in the *ApiAP2* G4mut version, nor in the *rifin* transcripts.

Both versions of both transcripts were used to explore the effect of an rG4 on *in vitro* protein translation. Prior to translation, the transcripts were subjected to various heat-denaturing and cation treatments, ensuring that rG4s would fold (when RNA was denatured then refolded in K^+^), or would not fold (e.g. when RNA was refolded in Li^+^). All transcripts were then translated in rabbit reticulocyte lysate. The HiBiT nanoluciferase fused to each protein provided a quantitative readout of protein production. Luminescence levels obtained from the *ApiAP2* genes showed that an rG4 could inhibit translation: protein production was two-fold lower from WT transcript than G4mut transcript, but only in rG4-forming conditions (refolded in K^+^) (Figure 4D, ANOVA, *p*-value = 0.0001). This was the only condition where a difference in protein production could be linked to the formation of an rG4. (Across the experiment there were some other differences in translation after different folding conditions (Supplementary Table 3) but these differences were smaller and could not be attributed to the presence or absence of an rG4: they were likely due to variation in other RNA secondary structures.) Finally, the same experiment performed on the *rifin* transcript, which evidently lacked an rG4, confirmed that subjecting these transcripts in G4-forming conditions had no impact on translation (Figure 4E; Supplementary Table 3).

### Ribosome profiling suggests that RNA G-quadruplexes can affect gene translation throughout the *P. falciparum* transcriptome

The effect of rG4s upon translation in *P. falciparum* was next investigated in the *in vivo* cellular context. Ribosome profiling and parallel RNA-Seq has been conducted in erythrocytic parasites at various stages (rings, trophozoites, schizonts and merozoites), enabling calculation of ribosome density per messenger RNA, or ‘translation efficiency’ (TE) ^27,28^. We calculated TE of all genes found to contain folded rG4s and compared this with the TE of all other genes in the transcriptome that were expressed at the same stage (Figure 5A,B). In rings, there was no significant difference; in trophozoites, TE was higher on average in rG4-containing genes, whereas in schizonts it was lower in rG4-containing genes. There was no clear relationship between TE and the number of rG4 motifs within a gene (data not shown).

**Figure 5:**
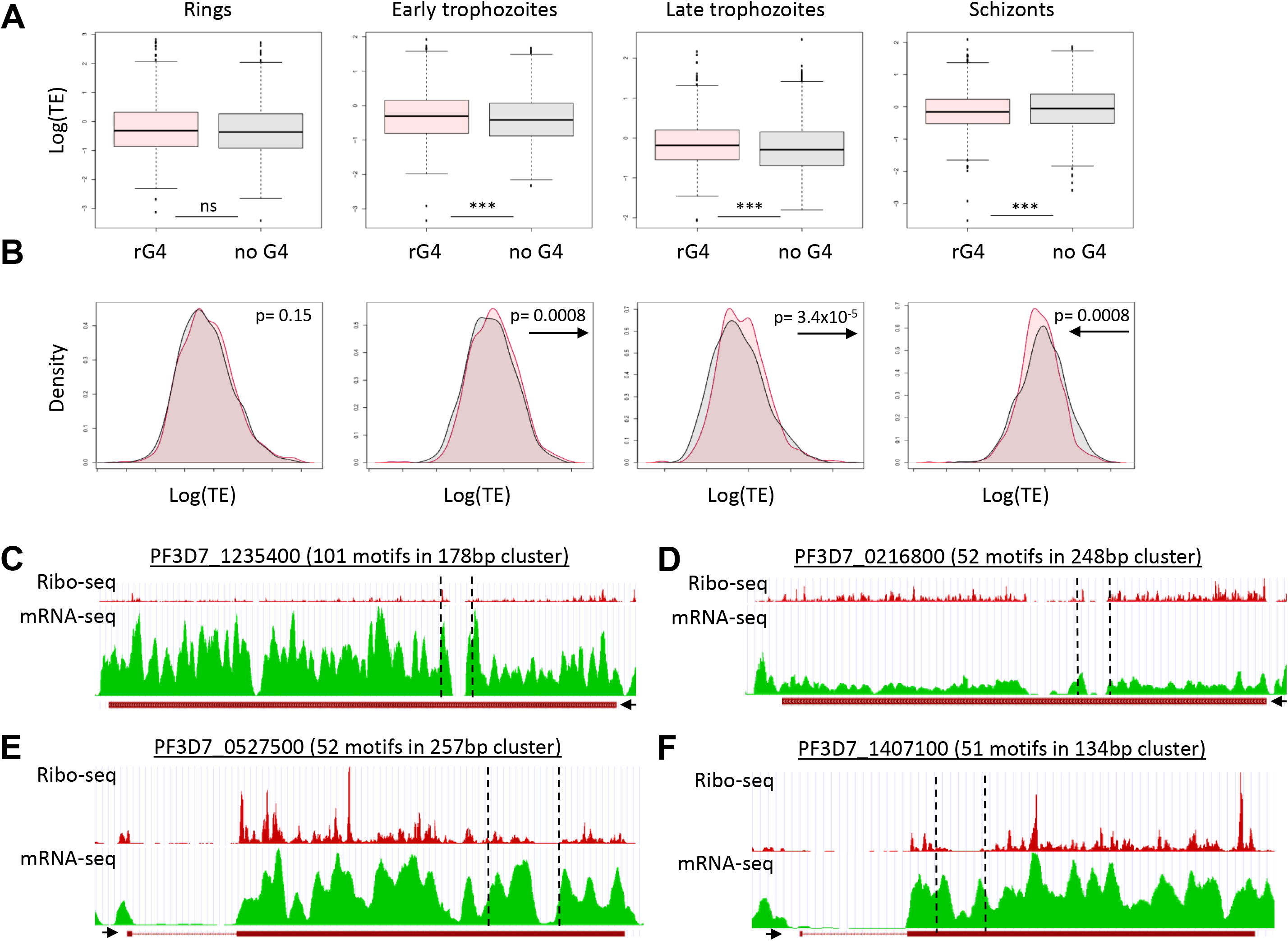
Translation efficiency of rG4-encoding genes. (A, B) Translation efficiency of all expressed genes in the transcriptome at four stages of the erythrocytic lifecycle, based on data from (28), in the form of either box-and-whisker plots showing mean and interquartile range of Log(TE) (A) or histograms of Log(TE) values (B). Genes with *bona fide* rG4s (detected by rG4-Seq) are in red while genes predicted to be rG4-null are in blue. p-values were calculated using two-tailed Student’s T-test: ***, p<0.001. Arrows accompanying the p-values show the direction of the shift in TE for rG4-genes. (C-F) Profiles of Ribo-Seq coverage and mRNA-Seq coverage for four genes containing large clusters of rG4s. Data obtained from ^28^, visualised via the GWIPS-viz Genome Browser ^50^. The locations of the rG4 clusters are marked with dotted lines. The 5’-3’ direction of the gene is indicated with a black arrow.

This is a transcriptome-wide analysis: it shows the average translational efficiency across hundreds of genes with very different characteristics. Therefore, we also investigated the individual profiles of a few genes encoding large clusters of overlapping rG4 motifs, to detect any relationship between the location of an rG4 cluster and the stalling or loss of elongating ribosomes. Five genes were selected that had the largest clusters of rG4s in which folding was detected, and four out of these five genes had convincing evidence of blood-stage expression in RNA-Seq. Figures 5C-F show the RNA-Seq and Ribo-Seq profiles for these four genes. Both types of coverage tended to drop in the region of the rG4 cluster, making it difficult to discern whether ribosomal occupancy dropped over and above the apparent drop in sequencing coverage. However, in at least one instance (Figure 5F), the loss of Ribo-Seq coverage clearly exceeded the loss of RNA-Seq coverage, and in two cases (Figure 5C, E) the overall Ribo-Seq coverage downstream of the cluster dropped, suggesting a reduction in translating ribosomes. Importantly, the inability to sequence these regions efficiently does, in itself, suggest the presence of G-quadruplexes: either in the RNA, where they may cause fragmentation bias in poorly-denatured transcripts, or in the sequenced cDNA, where they may cause DNA polymerase stalling.

### Reporter genes demonstrate that RNA G-quadruplexes can affect the translation of *P. falciparum* genes *in vivo*

Finally, we investigated the translation of individual rG4 reporter genes *in vivo* using cultured parasites. For this experiment, we excluded genes with large clusters of rG4s, such as those in Figure 5, because these would be difficult to manipulate experimentally. Instead we focussed on genes with a single detected rG4 that could be point-mutated to remove the rG4 without altering the protein sequence. This approach is shown in Figure 4A for the *in-vitro*-translated *ApiAP2* gene. A similar approach was taken with two additional genes, one gene with a long-loop 3-quartet rG4 encoding a rifin protein (PF3D7_0700200) and the other gene with a 2-quartet rG4 encoding a putative homolog of the DNA repair protein RAD54 (PF3D7_0803400) (Figure 6A).

**Figure 6:**
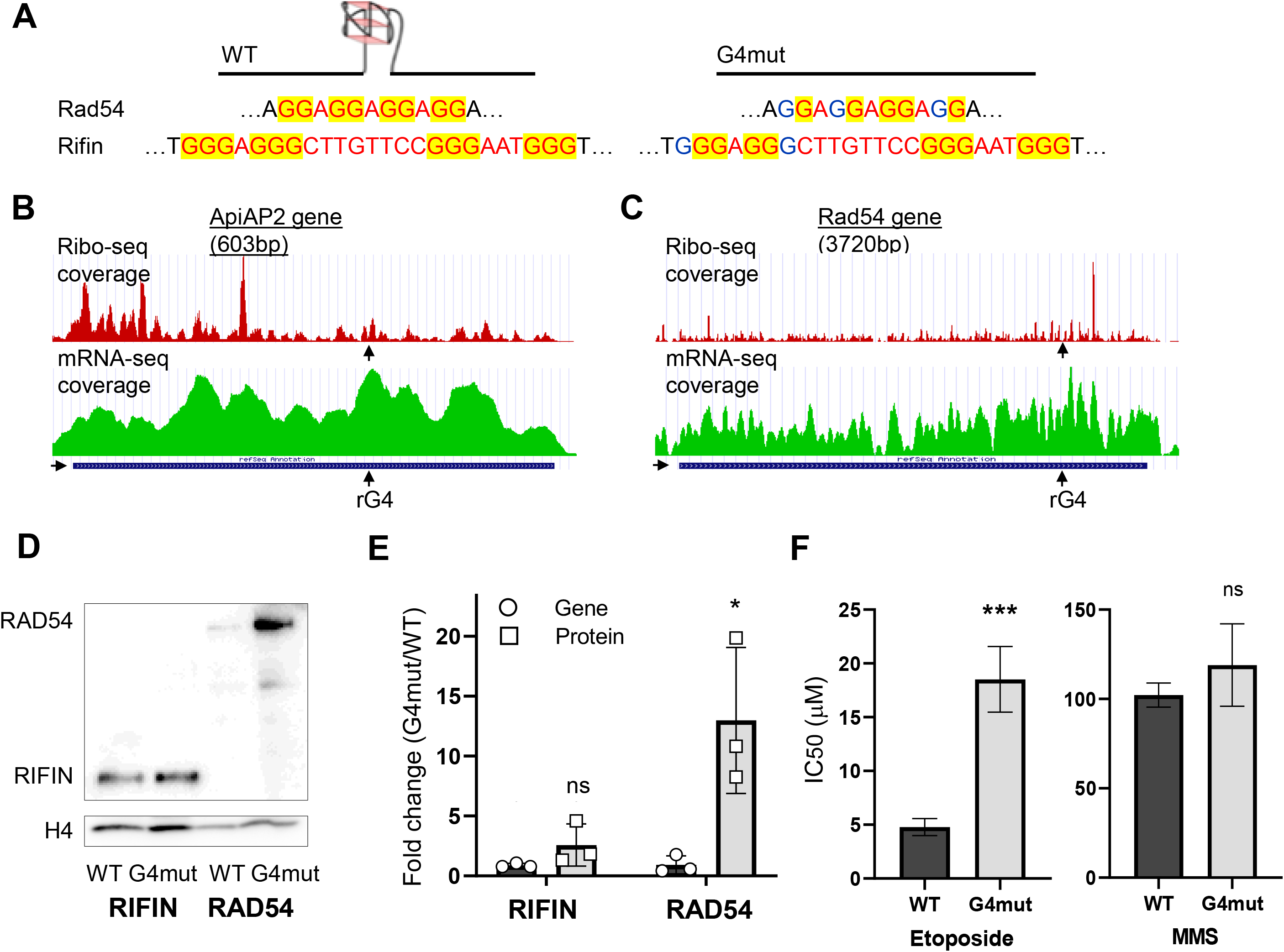
Translation of rG4-encoding reporter genes *in vivo*. (A) Schematic showing the WT and G4mut forms of the *Rad54 and rifin* genes with the rG4-forming motifs intact or mutated. (B,C) Profiles of Ribo-Seq coverage and mRNA-Seq for two of the selected rG4 reporter genes, *Rad54* and *AP2*. Data obtained from ^28^, visualised via the GWIPS-viz Genome Browser ^50^. The locations of the rG4s are marked. The 5’-3’ direction of the gene is indicated with a black arrow. (D) Western blot showing HA-tagged Rad54 and Rifin protein in the WT and G4mut reporter lines. Control blot shows histone H4 as a loading control. Image is representative of blots from three separate experiments. (E) Quantification by densitometry of the western blot data shown in (D) from three separate experiments, with the amount of HA-tagged protein controlled to histone H4. Quantification of the corresponding reporter-gene transcripts via qRT-PCR, controlled to the transcript levels of three housekeeping genes, is also shown. p-values are from two-tailed T-test: *, p<0.05, ns, not significant. (F) Comparison of DNA-damage sensitivity in the Rad54 WT and G4mut parasite lines, measured as 50% inhibitory concentrations of etoposide and MMS. Data are shown as mean and standard deviation of IC_50_ from four independent biological replicate experiments, each conducted in technical triplicate. p-values are from two-tailed T-test: ***, p<0.001, ns, not significant.

Two of these genes could be detected in the existing Ribo-Seq dataset (the *rifin* gene, which belongs to a large family, could not be uniquely identified) so their individual RNA-Seq vz Ribo-Seq coverage was examined. Changes in the region of the single rG4 were not clearly seen (Figure 6B, C), in contrast to the large drops in coverage within rG4 clusters that are shown in Figure 5. However, the *ApiAP2* gene did show a general drop in Ribo-Seq coverage towards the 3’ half of the gene, particularly in trophozoite stages (Supplementary Figure 3), occurring within ~150bp of the rG4 location (Figure 6B). The *RAD54* gene showed low Ribo-Seq coverage in general, even in the trophozoite stages when the mRNA was most abundant (Supplementary Figure 3), but there was marked accumulation of ribosomes within ~200bp of the rG4 (Figure 6C). These data suggested that an rG4 might affect the translation of these transcripts by promoting stalling or loss of translating polysomes, but to confirm this, a controlled *in vivo* experiment was clearly required.

We attempted to modify all three reporter genes in the parasite genome to carry a 3’ 3xHA tag. The rG4 was either mutated or left intact, generating paired parasite lines with ‘WT’ or ‘G4mut’ tagged genes. Two of these three pairs were successfully generated in parasites: the *RAD54* gene (PF3D7_0803400) and the *rifin* gene (PF3D7_0700200). Transcription of each reporter gene was measured by qRT-PCR at the erythrocytic stage of maximal transcription, controlled to the average transcriptional level of three housekeeping genes. Translation was simultaneously assessed by western blotting for the HA tag, using histone H4 as a loading control (Figure 6D). At the transcriptional level, neither reporter gene was strongly affected by the presence/absence of the rG4 (as expected in the case of the 2-quartet G4s, which are likely to be stable only in RNA, not DNA). At the translational level, however, the *RAD54* gene was clearly affected, with protein levels being an order of magnitude higher when the rG4 was mutated (p = 0.027, n=3). Translation of the G4mut *rifin* gene also trended upwards, but failed to reach statistical significance (p = 0.082, n=3) (Figure 6E). To examine whether the altered levels of RAD54 protein influenced parasite biology, we tested the sensitivity of the WT and G4mut parasite lines to DNA damage, because RAD54 is expected to play a role in the repair of DNA double strand breaks (DSBs). We therefore treated parasites with a DSB-inducing agent, etoposide, and also a DNA alkylating agent which is not a direct inducer of DSBs. The G4mut line, which expresses extra RAD54, was specifically less sensitive to etoposide when compared to the WT line (Figure 6F).

The strong biological effect of the presence/absence of an rG4 in the *RAD54* gene led us to examine whether this variation could occur in wild strains of *P. falciparum*. Hundreds of such strains have been sequenced from across the endemic world and indeed rG4-disrupting mutations do exist: a G-to-T mutation in the first guanine of the motif (<u>TG</u>A<u>GG</u>A<u>GG</u>A<u>GG</u>) appears commonly in 74 strains recorded in PlasmoDB ^29^, while a G-to-A mutation in the final guanine has also been recorded in a single Gambian strain.

## DISCUSSION

This study is the first experimental investigation of rG4s in the malaria parasite *P. falciparum*, an early-diverging prokaryote with a highly A/T-biased genome and many unusual features in its transcriptome ^17,30^. We demonstrate that rG4s can be found in this parasite and can affect gene translation.

Sequences with the potential to form G-quadruplexes are generally scarce in the *P. falciparum* genome and detection of rG4s using structure-specific antibodies or dyes is unlikely to give a full picture because such reagents probably do not detect rG4s at single-motif resolution ^6^. Furthermore, rG4s are likely to be dispersed throughout the cytoplasm, not arranged in arrays like telomeric DNA quadruplexes, so their visual detection is a particular challenge, even in large human cells ^24^. Accordingly, we found limited microscopic evidence for rG4s in *P. falciparum*.

By contrast, rG4-seq has very high resolution and it did reveal a surprisingly large number of rG4s in *P. falciparum*. Nevertheless, this method also has limitations. The detection of low-abundance transcripts is inevitably limited by low sequencing coverage, and in *P. falciparum* the detection of rG4s in UTRs was particularly limited because these UTRs are extremely A/T-rich, repetitive and resistant to NGS sequencing and assembly. Hence, even accepting the very low guanine density of *P. falciparum* UTRs, these rG4s were almost certainly under-detected (just 149 of 2569 motifs were in UTRs, a stark contrast with the high rG4 density found in human and plant UTRs ^18,19^). In fact, the detection of rG4s in all repetitive regions of the *P. falciparum* transcriptome proved to be limited by assembly issues. In future, long-read sequencing technology could be used to ameliorate this.

Our dataset contained a striking preponderance of non-canonical rG4 motifs, similar to the bias reported in the human rG4-ome ^18^, although *P. falciparum* showed an even higher proportion of ‘degenerate’ guanine-rich motifs. rG4 clustering was also a common feature in both human and *Plasmodium* genes: this may ensure that the biological effects of these motifs are robust to point mutations. It has also been proposed that biologically-important G4s may maintain ‘spare tyres’ of additional guanine tracts, making them more robust under oxidative stress which can oxidise guanine residues and thus disrupt G4s ^31^. *Plasmodium* parasites experience high levels of oxidative stress due to their endogenous metabolism and inflammatory responses in the host. Overall, it seems likely that both the existence and the clustering of rG4-forming motifs in some *P. falciparum* genes may be under positive evolutionary selection.

Another factor contributing to rG4 clustering is the interesting association that we observed between rG4s and sequences encoding low complexity peptide repeats, which are extremely common in *P. falciparum* ^32^. These LCRs can constitute broad regions that are relatively high in G/C due to the usage of G/C-rich codons, and they therefore frequently contained clusters of rG4 motifs, whereas non-LCR rG4s tended to be isolated and G/C-rich only within the motif itself. It remains unclear why LCRs are so heavily maintained in *P. falciparum* proteins but at least some of them are probably functional, for example in immune avoidance ^32–34^, and it is likely that a large subset of the rG4s in this transcriptome are a ‘byproduct’ of evolutionary selection for LCRs. It would be particularly interesting if the converse was also true: that selection for clusters of rG4s in certain genes has led to the maintenance of LCRs.

In seeking biological functions for rG4s in *P. falciparum* genes, we found several classes of genes that were enriched for rG4s, including variantly-expressed virulence genes, stress-response genes and genes encoding chromatin or DNA-modifying proteins. The rG4s in *var* genes were probably actually under-detected, both due to the repetitive nature of many of these motifs, and because the *var* gene family is mostly transcriptionally silenced. Reasons for DNA quadruplexes being associated with *var* genes have been proposed ^4,14^, but it remains unclear if rG4s have additional roles in post-transcriptional *var* gene regulation. Why genes for DNA-binding proteins should be enriched for rG4s is also unknown, but this feature appears in the human rG4-ome too ^18^. These essential genes may require particularly close control with layers of post-transcriptional regulation, since their transcripts are known to be generally enriched in conserved RNA structures ^35^. Finally, for stress-responsive genes it would make sense to evolve several layers of regulation in a parasite that experiences regular stresses from host immune responses and from temperature and environmental changes accompanying transitions between hosts. In plants, the transcripts of stress-responsive genes have been reported to be particularly unstructured and prone to degradation after severe heatshock ^36^; perhaps in *P. falciparum* it could instead be beneficial to stabilise these transcripts via stable quadruplex structures.

Experimentally, we have reported here the first examples of rG4-dependent gene translation in *P. falciparum*. Our results conform to a common model in which sense-strand rG4s can inhibit ribosomal translation: this effect appeared in the *AP2* reporter gene under *in vitro* translation and also in two reporter genes *in vivo*. (Notably, the effects of rG4s can be modulated by binding proteins so they are not always the same *in vitro* and *in vivo*. There are some examples of transcriptional enhancement by rG4s *in vivo* ^37^, although inhibition is probably more common, as was reported recently in a transcriptome-wide study in plants ^19^. Our transcriptome-wide analysis of translation efficiency in *Plasmodium* suggests a stage-specific effect, with rG4s affecting translation differently in trophozoite-expressed versus schizont-expressed genes.)

We detected a particularly strong translation-inhibiting effect from the 2-quartet rG4 in the *Rad54* reporter gene, and a weaker effect in the *rifin* gene. This may be partly because the rG4 was completely ablated in the G4mut version of *Rad54*, whereas the G4mut *rifin* gene retained a potential 2-quartet rG4 instead of a 3-quartet motif. Of note, the rG4 in the *Rad54* gene also overlaps with a short stretch of low complexity peptide sequence (DDDDDEEEEE), in which mutagenesis to create the G4mut version altered three GAG codons to GAA. *P. falciparum* has a strong codon preference for GAA over GAG ^38^, so this change, besides the rG4 disruption, could also have contributed to increased translation efficiency. The effect in the *Rad54* gene was particularly interesting because it is likely to be replicated in wild strains of *P. falciparum*, in which the rG4-forming motif is commonly mutated, and because it affected the DNA repair capacity of mutant parasites., This could in turn influence important factors like genomic mutation rates and responses to DNA-damaging antimalarial drugs such as artemisinins.

In conclusion, the G-quadruplex biology of malaria parasites remains an exciting area for future study. This is only the first report on the effects of rG4s in *P. falciparum* and much of the biology associated with these interesting motifs probably remains to be discovered.

## MATERIALS AND METHODS

### Parasite culture and drugs

The 3D7 strain of *P. falciparum* was obtained from the Malaria Research and Reference Reagent Resource Center (MR4). Parasites were cultured as previously described ^39^, in gassed chambers at 1% O2, 3% CO2, and 96% N2. Parasite growth and morphology was assessed on blood smears stained with Hemacolor (Merck). Synchronised parasite cultures were obtained by performing two treatments with 5 % D-sorbitol ^40^ 42 h apart, to yield an approximately 4-h window of ring stage parasites.

### Visualisation of rG4s by fluorescent microscopy

Air-dried slides of 3D7 parasites were made, after saponin-mediated release of the parasites from host erythrocytes. Cells were fixed with 8% formaldehyde/HEPES pH 7.4 for 10 minutes and then with 4% formaldehyde/HEPES pH 7.4 for 10 minutes at room temperature (RT). Permeabilisation was done in 0.05% TritonX for 10 minutes. Enzymatic treatments were performed as followed: 0.12 U μl^−1^ Turbo DNase (Albion) or 5 U RNase A and 20 U RNase T1 (RNase Cocktail enzyme mix, Thermo Fisher Scientific) for 1 h at 37°C. Slides were washed three times for 5 minutes in PBS, adding 2 μg/mL DAPI (4’,6-diamidino-2-phenylindole; Thermo Fisher Scientific) and 1 μM QUMA-1 ^23^ to the final wash, followed by three washes with DEPC treated water. Slides were mounted using ProLong diamond antifade (Thermo Fischer Scientific). Visualisation was performed on a Nikon SA microphot microscope.

### Preparation of polyA RNA

RNA was harvested from two independent parasite cultures using Trizol (Invitrogen), as previously described ^41^. RNA was treated with DNase1 for 30 minutes and the absence of DNA was confirmed via PCR across an intronised gene, as previously described ^42^. RNA was then subjected to polyA purification via the NEBNext Poly(A) mRNA Magnetic Isolation Module (NEB), and quantified using a Nanospec 1000 (Thermoscientific, USA).

### rG4-seq library preparation

The rG4-seq libraries were prepared as per our previous study, with minor modifications ^18^. Approximately 1μg of mRNA was obtained after polyA RNA enrichment. Random RNA fragmentation was performed in buffer (final 1X: 40 mM Tris-HCl pH 8.0, 100 mM LiCl, 30 mM MgCl_2_) at 95 °C for 45 s to result in RNA fragment size of ~250 nt, followed by RNA clean and concentrator-5 (Zymo research). Next, 3′ dephosphorylation was performed by using 8 μl RNA sample, 1 μl 10× T4 PNK buffer and 1 μl T4 PNK enzyme (NEB) at 37 °C for 30 min. Then, 3′ adapter ligation was conducted by adding 10 μl sample from above, 1 μl of 10 μM 3′ rApp adaptor (5′-/5rApp/AGATCGGAAGAGCACACGTCTG/3SpC3/-3′), 1 μl 10x T4 RNA ligase buffer, 7 μl PEG8000 and 1 μl T4 RNA ligase 2 K227Q (NEB) at 25 °C for 1 h, followed by RNA clean and concentrator and eluted in nuclease-free water. The eluted sample was then split into two parts for 150 mM Li^+^ and 150 mM K^+^ for reverse transcription (~12 μl each), with 1 μl of 5 μM reverse primer (5′-CAGACGTGTGCTCTTCCGATCT-3′) and 6 μl of 5x reverse transcription buffer (final concentration 20 mM Tris, pH 7.5, 4 mM MgCl_2_, 1 mM DTT, 0.5 mM dNTPs, 150 mM LiCl or 150 mM KCl). The mixture was heated at 95 °C for 1.5 min and cooled at 4 °C for 1.5 min, followed by 37 °C for 15 min before 1 μl of Superscript III (200 U/μL) was added. The reverse transcription was carried out at 37 °C for 40 min, followed by treatment of 1 μl of 2M NaOH at 95 °C for 10 min. 5 μl of 1M Tris-HCl (pH 7.5) was added to neutralize the solution, before the sample was cleaned up by RNA clean and concentrator and eluted in nuclease-free water. To the purified and eluted cDNA samples (8 μl), 1 μl of 40 μM 5′ adapter was added (5′/5Phos/AGATCGGAAGAGCGTCGTGTAGCTCTTCCGATCTNNNNNN/3SpC3/3′). The sample was heated at 95 °C for 3 min, cooled to room temperature, and 10 μl of 2x Quick T4 ligase buffer and 1 μl Quick T4 DNA ligase (NEB) were added and incubated at room temperature overnight. The ligated cDNAs were purified by pre-cast 10% urea denaturing TBE gel and the size 90-450 nt was cut, followed by the gel extraction step using crush and soak methods. Next, a PCR reaction (20 μl) was performed using 95 °C: 3 min, 12 cycles of each temperature step (98 °C: 20 s, 65 °C: 15 s, 72 °C: 40 s); 72 °C: 1 min, 1 μl 10 μM forward primer (5′ AATGATACGGCGACCACCGAGATCTACACTCTTTCCCTACACGACGCTCTTCCGATCT 3′) and 1 μl 10 μM reverse primer (e.g., index 2) (5′ CAAG CAGAAGACGGCATACGAGATACATCGGTGACTGGAGTTCAGACGTGTGCTCTTCCGATCT 3′), 8 μl DNA template and 10 μl 2x KAPA HiFi readymix. The amplified libraries were purified with 1.8% agarose gel for 50 mins at 120V, and the size 150-400 bp was sliced and extracted with Zymoclean Gel DNA Recovery Kit. The purified libraries underwent qPCR with KAPA Universal Quant Kit and were subjected for next-generation sequencing on NEXTseq 500 (Illumina).

### rG4-seq data analysis

Pre-processing and short read alignment of rG4-seq sequencing data were conducted as previously described ^43^. Reverse transcriptase stalling (RTS) site analysis and rG4 calling were conducted using the rG4-seeker pipeline ^43^. The definition of rG4 structural motifs has been previously described ^18,43^. The intersection and union of detected rG4s across the four rG4-seq experiments (2 replicates, 2 conditions) were computed based on the overlapping of genomic coordinates of the rG4 sequences.

### *P. falciparum* genome sequence and transcriptome annotation sources

The reference genome and transcriptome annotation for *P. falciparum* (3D7) were downloaded from PlasmoDB ^29^. As the transcriptome annotation from PlasmoDB lacked UTR information, the 2000 nt upstream and downstream of each annotated protein coding gene were flagged as ‘predicted’ 5’UTR/3’UTR regions, as described in ^4^. The curated UTR annotations reported by Chappell *et al.* ^44^ were then appended to the annotation. UTR regions that were encompassed by both predicted and curated UTRs were considered as curated UTRs.

### Annotation of repeats in the *P. falciparum* genome

DNA repetitive elements in the *P. falciparum* genome were annotated using RepeatModeler and RepeatMasker software at default settings ^45^. The overlap between rG4 sequences and repetitive elements was computed using bedtools ^46^.

### G/C content calculation

G/C content of rG4 sequences and rG4 flanking sequences was calculated directly. The G/C content of transcripts was calculated in sliding windows of 50 nt and at increments of 10 nt. The statistical significance of differences in G/C content for various set of sequences was computed using Wilcoxon rank-sum test for two samples.

### Low-complexity peptide region (LCR) analysis

The peptide sequence matching an rG4 and its ±50 nt flanking sequence was first extracted from the full protein sequence of the rG4-harbouring gene. Any peptide sub-sequence of at least 10 amino acids that matched the following composition bias or tandem-repeat definition was then exhaustively searched for:

Composition bias – 80% of the peptide sequence is composed of 1 kind of amino acid residue; or 100% of the peptide sequence is composed of 2 kinds of amino acid residue, where one kind takes <80% and the other kind takes >20%.

Tandem repeats – the peptide sequence is composed of at least 3 repeat units of 3-10 amino acids. Imperfections in the repeats (substitution/insertion/deletion of 1 amino acid residue) were tolerated if the number of imperfections was ≤ 2 (for peptide sequences longer than 11 residues) or ≤ 1 (for other peptide sequence lengths).

If more than one sequence or type of LCR was identified, the longest LCR with least number of imperfections was chosen for reporting. If both composition bias and tandem repeats were present, the composition bias was chosen for reporting.

### Gene Ontology (GO) analysis

GO term annotations for *P. falciparum* genes were downloaded from PlasmoDB ^29^. GO enrichment analysis were conducted on the PANTHER platform ^47^, using a de-duplicated list of rG4-harbouring gene IDs as input.

### *In vitro* transcription and translation

The genes *ApiAP2* (PF3D7_0934400) and *rifin* (PF3D7_1254400) were PCR amplified using Q5 DNA polymerase (NEB) from *P. falciparum* cDNA with overhang primers to insert a 5’ *Mlu*I and a 3’ *Sma*I restriction site. After digestion of the PCR product using *Mlu*I and *Sma*I for 1 h at 37°C, the DNA fragments were cloned into a version of the pTnT vector (Promega) where the multiple cloning site had been replaced by a synthetic construct (Genewiz) composed of a 5’ HiBiT tag (Promega), a new multiple cloning site and 3’ 3xHA tag. Disruption of the G4 motif in both genes was performed by site-directed mutagenesis using Pfu Turbo (Agilent) and *Dpn*I (ThermoFisher). Prior to *in vitro* transcription, each construct was linearised using *BamH*I (ThermoFisher) and purified via phenol/chloroform and ethanol precipitation. 1 μg of linearised plasmid was incubated for 2 h at 37°C with 1U T7 RNA polymerase (ThermoFisher), 5 mM DTT, 4 nM NTPs (ThermoFisher) and 50U Superase (Invitrogen). Transcription products were treated with 10 Kunitz units of RNase-free DNase (ThermoFisher) at 37°C for 45 minutes, then purified as above.

Translation was performed in Flexi® Rabbit Reticulocyte Lysate (Promega) supplemented with amino acid mixture minus methionine and leucine, 60nM each, Complete Protease inhibitor (Roche) and 20U Superase protease inhibitor (Invitrogen). Transcripts were first denatured and refolded by incubation at 95°C for 5 minutes followed by 15 minutes at RT with 150 mM KCl or LiCl. Translation was performed for 60 minutes at 30°C with 300 ng of transcript in three independent reactions. Each reaction was stopped by the addition of 4:1 (v:v) blocking solution (1 mg of cycloheximide, Complete protease inhibitor in PBS). Luciferase activity was measured via the HiBiT tag using Nano-Glo® HiBiT Extracellular Detection System (Promega) on a FLUOstar Omega (BMG Labtech). Data were analysed with Graphpad Prism using a two-way ANOVA and a Bonferroni correction for multiple testing.

### NMM and QUMA-1 ligand-enhanced fluorescence assay

Enhanced fluorescence assays with both ligands – NMM and QUMA-1 – were performed as previously reported ^48^. In summary, in a reaction volume of 100 μL, 1 μM RNA was mixed with 10 mM LiCac (pH 7.0) and 150 mM KCl or LiCl. Samples were annealed at 95°C for 5 minutes and cooled to RT for 15 minutes for renaturation to occur. Samples were then transferred into a 1 cm path-length quartz cuvette and 5 μL of 20 μM ligand, NMM or QUMA-1, (final concentration of 1 μM) was then added. Excitation was at 394 nm and 555 nm for NMM and QUMA-1 respectively, and the emission spectrum was collected from 550-750 nm and 575-800 nm respectively. Measurements were performed using a HORIBA FluorMax-4 and spectra were acquired every 2 nm at 25°C for both WT and rG4mut RNA oligos. For full transcripts rather than RNA oligos, the same procedure was followed after *in vitro* transcription, with a few alterations: 200nM transcript was used with 4 μM QUMA-1. Spectra were measured using a Cary Eclipse Fluorescence Spectrophotometer (Agilent).

### Circular Dichroism assay

As previously reported ^48,49^, reactions were set up in 2 mL samples of 5 μM RNA prepared in 10 mM LiCac (pH 7.0) and 150 mM KCl or LiCl. All RNAs were folded as described above and measurements were conducted using a Jasco CD J1500 spectropolarimeter and a 1 cm path-length quartz cuvette. Spectra were acquired every 1 nm from 220 to 310 nm at 25°C for WT and rG4mut RNA oligos. All spectra reported are the average of 2 scans with a response time of 2 s/nm, normalised to molar residue ellipticity and smoothed over 5 nm. All data were analysed with Spectra ManagerTM suite (Jasco Software).

### Thermal melting monitored by UV spectroscopy

Thermal melting monitored by UV spectrocopy was performed using a Cary 100 UV-Vis spectrophotometer and a 1 cm path-length quartz cuvette with a reaction volume of 2 mL ^49^. Samples were prepared and renatured as per the circular dichroism experiment above. Data were collected over 0.2°C increments while heating over the temperature range 5-95°C. The unfolding transitions were monitored at 295 nm for the WT and rG4mut RNA oligos to look for the inverse melting G4 signature. Data were smoothed over 5 nm.

### Translation efficiency analysis

Translation efficiency was calculated as the ratio of ribosome-protected footprints (Ribo-Seq) to mRNA reads (RNA-Seq) across each mRNA, using published *P. falciparum* Ribo-Seq data ^28^. Significant differences between the TE distributions of genes harbouring or lacking rG4s were assessed by Student’s t-test. Coverage of Ribo-Seq and RNA-Seq for individual genes was visualised via GWIPS-viz Genome Browser (https://gwips.ucc.ie, ^50^) or the Mochiview browser ^51^.

### Plasmid construction and parasite transfection

To generate the rG4 reporter parasite lines, 3 genes harbouring an rG4 on the positive strand were selected, *rifin* (PF3D7_0700200), *RAD54* (PF3D7_ 0803400) and *AP2* (PF3D7_0934400). The 3’ portion of each gene containing the rG4 was cloned in-frame with the 3xHA tag in a pSLI vector designed for selectable integration into the *P. falciparum* genome ^52^. Briefly, all PCR reactions were performed using Phusion (NEB) with primers to add a 5’ *Eco*52I restriction site and a 3’ *Kpn*I restriction site (see Supp Table 3 for oligos). PCR products and plasmids were digested with *Eco*52I and *Kpn*I (Thermo Fisher Scientific), purified using QIAquick PCR Purification Kit (Qiagen), then cloned using T4 DNA ligase (Thermo Fisher Scientific). The G4mut variants with point-mutations designed to disrupt the G4 motif were generated as previously described in the methods section on *in vitro* transcription. All plasmids were transformed into PMC103 electro-competent bacteria using a Gene Pulser Xcell Electroporation System (Bio Rad) and 0.1cm cuvettes (Bio Rad). All constructs were checked by restriction digestion and sequencing.

Plasmids were transfected into the 3D7 strain of *P. falciparum* using standard procedures ^53^. Transfectants were selected with 4 nM WR99210 (Jacobus Pharmaceuticals), then selection for parasites with genomic integration of the pSLI plasmid was performed with 400 μg.mL^−1^ of G418 (Sigma). Recombinant parasites were checked for correct rearrangement by PCR (Supp Table 3).

### Reporter gene expression analysis and protein quantification

All reporter parasite lines were synchronised twice with 5% sorbitol and total RNA and protein were then extracted at the lifecycle stage of peak gene expression ^29^. *RAD54* and *rifin* lines were harvested at 40hpi and AP2 at 48hpi.

After saponin lysis to release parasites from erythrocytes, total RNA was extracted using the RNeasy Mini Kit (Qiagen) according to manufacturer’s instructions. DNA contamination was removed by incubating the RNA extracts with 2 U TURBO DNase (Ambion) for 30min at 37°C. cDNA preparation was performed using SensiFAST cDNA Synthesis Kit (Bioline) and real-time qPCR was performed using SensiFast SYBR Lo-ROX kit (Bioline). RT-qPCR was carried using primer sets for the three reporter genes and three housekeeping genes (actin I (p100, PF3D7_1246200), serine tRNA ligase (p60, PF3D7_0717700), fructose-bisphosphate aldolase (p61, PF3D7_1444800). ΔΔCt analysis (Ct, threshold cycle) was used to calculate the relative copy number of each target gene relative to the average Ct of the three control genes. All experiments were conducted in three biological replicates, assessed in technical triplicate.

Secondly, parasite fractions for western blotting were prepared in parallel as previously described ^54^. Samples were loaded onto 4-12% polyacrylamide gels (BioRad) and electrophoresed at 100V for 60 minutes. Electrophoretic transfer to nitrocellulose membrane was carried out at 100V for 60 mins. Membranes were blocked in TBST with 5% milk protein and probed with the following antibodies: 1:1000 anti-HA (Roche), then 1:1500 goat anti-rat IgG-HRP (Dako); anti-histone H4 (Abcam), then 1:1000 goat anti-rabbit IgG-HRP (Abcam). Membranes were washed for 3 × 5 minutes in TBST after each antibody step. SuperSignal West Pico PLUS substrate (Thermo Fisher Scientific) was added for 1 minute and blots were imaged using a FluorChemM chemiluminescent detection camera (ProteinSimple).

### DNA damage sensitivity assay

The Malaria SYBR Green I-based fluorescence (MSF) assay was used to measure parasite growth in the presence of DNA damaging agents, essentially as previously described ^55^. Trophozoite-stage cultures of each parasite line were seeded in triplicate into 96-well plates at 1% parasitaemia, 4% haematocrit, with serial dilutions of the drugs etoposide (Millipore) or methyl methanesulphonate (MMS, Sigma). Plates were incubated for 48h in a gassed chamber at 37°C. Following this, 100 μL of sample from each well was transferred to a plate containing in each well 100 μL MSF lysis buffer (20 mM Tris pH 7.5, 5 mM EDTA, 0.008% saponin, 0.8% Triton X-100) supplemented with 0.2 μL/mL of SYBR Green I (Sigma). After a 1h incubation in the dark at room temperature, fluorescence was measured (excitation/emission 485/520 nm) in a FLUOstar Omega microplate reader (BMG Labtech). Percentage parasite growth was calculated as follows from the amount of DNA detected via SYBR Green I: 100x[μ_(s)_ - μ_(−)_/μ_(+)_ - μ_(−)_] where μ_(s)_, μ_(−)_ and μ_(+)_ are the means of the fluorescent readouts from sample wells (μ_(s)_), control wells with 100μM chloroquine (μ_(−)_, representing 0% growth), and control wells with non-drug-treated parasites (μ_(+)_, 100% growth). 50% inhibitory concentrations (IC_50_) for each drug on each parasite line were calculated using GraphPad Prism.

## DATA AVAILABILITY

Raw data from the rG4-seq experiment are available in SRA, accession number PRJNA706892. All other data are available within the manuscript and its supplementary figures.

## ACKNOWLEDGEMENT

We are grateful to Florian Noulin for early work on reporter gene design, to Tim Fitzmaurice (Cambridge Department of Physics) for the use of a fluorometer, to Harriet Mears (Cambridge Department of Pathology) for help with *in vitro* transcription and to Prof. Jia-heng Tan (Sun Yat-sen University) for the free gift of QUMA-1 for this study.

## FUNDING

This work was supported by the UK Medical Research Council [grants MR/K000535/1 and MR/L008823/1] to CJM. Shenzhen Basic Research Project [JCYJ20180507181642811], Research Grants Council of the Hong Kong SAR, China Projects [CityU 11101519, CityU 11100218, N_CityU110/17, CityU 21302317], Croucher Foundation [Project No. 9500030, 9509003], State Key Laboratory of Marine Pollution Director Discretionary Fund, City University of Hong Kong [projects 6000711, 7005503, 9680261] to CKK. A generous donation from Mr. and Mrs. Sunny Yang, the University Grants Committee Area of Excellence Scheme (AoE/M-403/16), and the Innovation and Technology Commission, Hong Kong Special Administrative Region Government to the State Key Laboratory of Agrobiotechnology (CUHK) to TFC. EYCC is supported by the Hong Kong PhD Fellowship Scheme. Armin Scheben copyedited this manuscript. Any opinions, findings, conclusions or recommendations expressed in this publication do not reflect the views of the Government of the Hong Kong Special Administrative Region or the Innovation and Technology Commission. The funders had no role in study design, data collection and interpretation, or the decision to submit the work for publication.

## AUTHOR CONTRIBUTIONS

FD conducted and designed experiments, analysed data and produced figures; EC and TFC wrote algorithms, analysed data, produced figures and wrote the manuscript; LMH prepared parasite RNA; CKK performed rG4-seq experiments; MIU conducted biophysical experiments; AJ conducted reporter-gene experiments; BC helped with *in vitro* translation experiments and analysis of translational efficiency; CJM and CKK designed the study, analysed data, produced figures and wrote the manuscript. All authors read and approved the final manuscript.

## CONFLICT OF INTEREST

The authors declare no conflict of interest.

## Notes

### Competing Interest Statement

The authors have declared no competing interest.

