## Supplementary material for "G-quadruplex RNA motifs influence gene expression in the malaria parasite *Plasmodium falciparum*": Supp figures and legends

### SUPPLEMENTARY FILES

Supp figure 1: **QUMA-1 staining of *P. falciparum* parasites**

Parasites were removed from their host cell and fixed in formaldehyde. Enzymatic treatments to selectively remove RNA or DNA were performed, before staining with QUMA-1 and DAPI.

Supp figure 2: **Biophysical analysis of the rG4-forming sequence from the AP2 reporter gene**

(A, B) Ligand enhanced fluorescence spectra showing fluorescence of rG4-encoding oligos (wild, wildtype sequence; Mut, rG4-mutated sequence) folded under $K^+$ as compared to $Li^+$ conditions in the presence of rG4 ligands QUMA-1 (A) and NMM (B).

(C, D) Circular dichroism performed on the same oligos in (A, B). Blue and red lines represent folding in $K^+$ and $Li^+$ ions. (C) CD spectrum of wild type sequence, showing a negative peak at ~240 nm and positive peak at ~262 nm, suggesting the formation of parallel topology. (D) CD spectrum of mutant sequence showing a lesser sign of rG4 formation in both $K^+$ and $Li^+$ conditions.

(E-H) UV Melting performed on the same oligos in (A, B). (E) UV melting of wild type oligo in $K^+$ condition, (F) UV melting of wild type oligo in $Li^+$ condition, (G) UV melting of mutant oligo in $K^+$ condition, (H) UV melting of mutant oligo in $Li^+$ condition. The observed Tm of the wild type is higher than the $Li^+$ condition, indicating the physiological stability of the rG4 under $K^+$ conditions.

Supp figure 3: **Translation efficiency of rG4-encoding reporter genes *in vivo* at 4 lifecycle stages**

Profiles of Ribo-Seq coverage and mRNA-Seq for two of the selected rG4 reporter genes, *AP2* (A) and *Rad54* (B), shown across lifecycle stages. Transcription of the *AP2* gene peaks in late trophozoites and schizonts; transcription of the Rad54 gene peaks in late trophozoites. Data obtained from [28], visualised via the Mochiview browser. The locations of the rG4s are marked.

Supp Table 1 **Gene-level summary of predicted $(G_3N_x)_4$ PQSs and their detection status in rG4-seq**

Supp Table 2 **Oligo sequences used in this study**

Supp Table 3 **Statistical analysis of *in vitro* translation data**

Detailed output of the 2-way ANOVA for the results of *in vitro* transcription of *AP2* and *Rif12* genes. A Bonferroni correction for multiple testing was applied. Statistics were obtained using GraphPad Prism.

Supp Data File 1 **List of 2,569 *P. falciparum* rG4s detected by rG4-seq**

Supp Data File 2 **Prediction outcome and repeat annotations of 335 $(G_3N_x)_4$ PQSs in *P. falciparum* protein coding genes**

Supp Figure 1

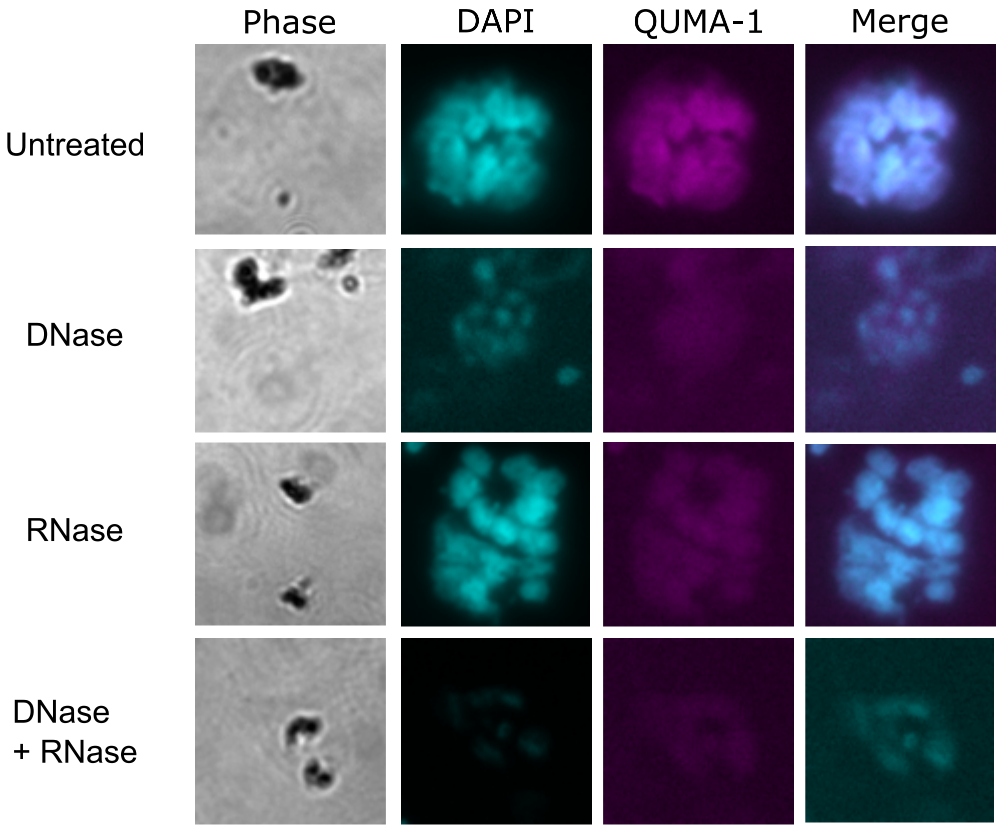


**QUMA-1 staining of *P. falciparum* parasites outside of the host cell**
Parasites were removed from their host cell and fixed in formaldehyde. Enzymatic treatment to selectively remove the different types of nucleic acids was performed before QUMA-1 staining.

Supp Figure 2

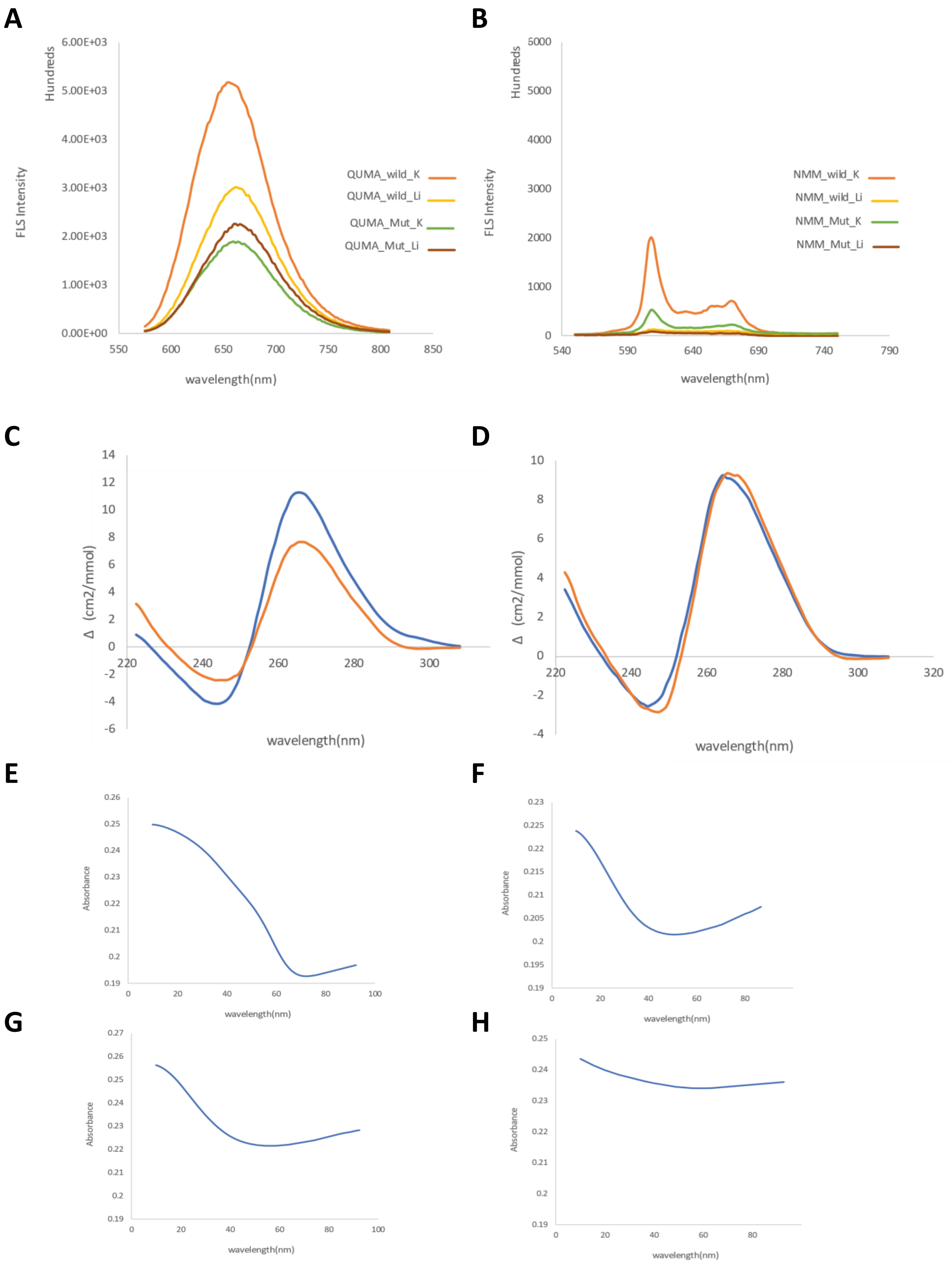

Supp Figure 3

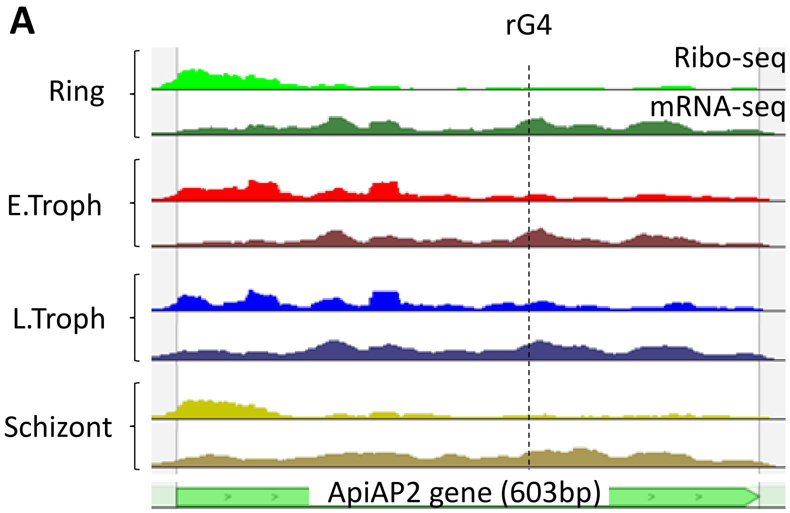


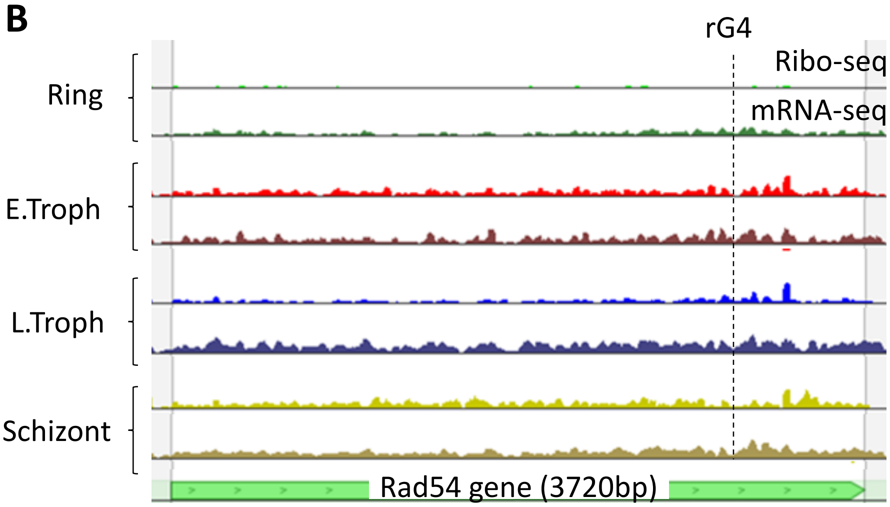
